# Electrophysiological Markers of Memory Consolidation in the Human Brain when Memories are Reactivated during Sleep

**DOI:** 10.1101/2022.06.08.495049

**Authors:** Jessica D. Creery, David J. Brang, Jason D. Arndt, Adrianna Bassard, Vernon L. Towle, James X. Tao, Shasha Wu, Sandra Rose, Peter C. Warnke, Naoum Issa, Ken A. Paller

**Affiliations:** Department of Psychology and Cognitive Neuroscience Program, Northwestern University, Evanston IL 60208; Department of Psychology, University of Michigan, Ann Arbor MI 48109; Department of Psychology, Middlebury College, Middlebury VT 05753; Department of Neurology, The University of Chicago, Chicago IL 60637; Department of Neurological Surgery, The University of Chicago, Chicago IL 60637

## Abstract

Human accomplishments depend on learning, and effective learning depends on consolidation. Consolidation is the process whereby new memories are gradually stored in an enduring way in the brain so that they can be available when needed. For factual or event knowledge, consolidation is thought to progress during sleep as well as during waking states, and to be mediated by interactions between hippocampal and neocortical networks. However, consolidation is difficult to observe directly, but rather is inferred through behavioral observations. Here, we investigated overnight memory change by measuring electrical activity in and near the hippocampus. Electroencephalographic (EEG) recordings were made in five patients from electrodes implanted to determine whether a surgical treatment could relieve their seizure disorders. One night, while each patient slept in a hospital monitoring room, we recorded electrophysiological responses to 10-20 specific sounds that were presented very quietly, to avoid arousal. Half of the sounds had been associated with objects and their precise spatial locations that patients learned before sleep. After sleep, we found systematic improvements in spatial recall, replicating prior results. We assume that when the sounds were presented during sleep, they reactivated and strengthened corresponding spatial memories. Notably, the sounds also elicited oscillatory intracranial EEG activity, including increases in theta, sigma, and gamma EEG bands. Gamma responses, in particular, were consistently associated with the degree of improvement in spatial memory exhibited after sleep. We thus conclude that this electrophysiological activity in the hippocampus and adjacent medial temporal cortex reflects sleep-based enhancement of memory storage.

**Significance Statement:** Sleep contributes to memory consolidation, we presume, because memories are replayed during sleep. Understanding this aspect of consolidation can help with optimizing normal learning in many contexts, and with treating memory disorders and other diseases. Here, we systematically manipulated sleep-based processing using targeted memory reactivation; brief sounds coupled with pre-sleep learning were quietly presented again during sleep, producing (a) recall improvements for specific spatial memories associated with those sounds, and (b) physiological responses in the sleep EEG. Neural activity in the hippocampus and adjacent medial temporal cortex was thus found in association with memory consolidation during sleep. These findings advance understanding of consolidation by linking beneficial memory changes during sleep to both memory reactivation and specific patterns of brain activity.

## Introduction

Memory research and sleep research have converged in recent years to yield increasingly strong evidence for the hypothesis that important memory processing takes place during sleep. A widely held view is that sleep is an ideal state for memory consolidation to occur, given reduced demand for sensory and executive function. Sleep may be particularly relevant for newly formed memories for facts and events (declarative memories) and probably for other types of memory as well (although here we emphasize declarative memories). Many investigators have argued that sleep-based consolidation involves a recapitulation of recent memory representations via conjoint operation of hippocampal and neocortical networks (1-5). The changes in memory storage that mediate consolidation would likely take place in these same networks. Yet, it has been difficult to fully characterize the neural operations that underlie consolidation.

The assumption that consolidation is tied to the reactivation of recently formed memories is reasonable, given that memory change would require some sort of activation of the existing memory. Yet, consolidation is more than just memory reactivation. Observations of memory reactivation are therefore an important part of this research but not sufficient; quantitative memory change must also be part of the empirical basis for understanding memory storage and consolidation.

Research on hippocampal activity in rodents has defined the phenomenon termed *hippocampal replay* (6) whereby patterns of place-cell firing first observed in a novel waking context are observed again during subsequent sleep (7, 8). Coordinated replay between the hippocampus and neocortical regions such as visual cortex (9) is suggestive of the communication across regions that might be pivotal for consolidation. Whether hippocampal replay occurs with a concurrent experience of conscious retrieval is unclear, as is whether it produces changes in memory storage. Memory consolidation during sleep may occur in many species, including songbirds (10).

Research on human sleep has produced evidence linking memory processing with deep stages of sleep in particular. Arguably the strongest such evidence has been obtained using the method of Targeted Memory Reactivation (TMR) whereby memory processing during sleep can be systematically altered. For example, Rasch and colleagues (11) administered a spatial learning task with a background rose odor, and then presented this odor again during sleep. Results showed that TMR with the same odor during sleep improved performance after sleep, consistent with the suggestion that the odor reactivated the spatial memories. This memory benefit was found with odor presentations during slow-wave sleep (SWS), but not during rapid eye-movement sleep (REM). Whereas the odor TMR cue in this study influenced all memories learned in the same context, Rudoy and colleagues (12) showed that auditory TMR cues could produce benefits for specific spatial memories. Before sleep, participants learned a set of unique object-location associations, each linked with a distinct sound, and then a subset of the sounds were played during SWS to reactivate corresponding memories. Additional studies of TMR have replicated and extended these findings with many sorts of learning (13, 14).

A reasonable hypothesis is thus that memory consolidation can progress by virtue of hippocampal-neocortical interactions during SWS in conjunction with the reactivation of recently formed memories. Findings from studies of amnesia have long supported the idea that these hippocampal-neocortical interactions are needed (15, 16). Hippocampal projections to entorhinal cortex, and from there to parahippocampal and perirhinal cortex, can presumably provoke widespread neocortical activity for specific memories. This activity could thereby produce a recapitulation of information principally represented in cortical networks — which specific networks would vary in accordance with which types of specialized cortical processing are needed for a given fact or event. In keeping with these ideas about hippocampal-neocortical interactions, Peigneux and colleagues (17) showed that cerebral blood flow in the hippocampus during slow-wave sleep was correlated with post-sleep performance in a route-learning task initially learned before sleep. Moreover, a subsequent study showed that sleep appeared to improve memory in association with increased functional connectivity between hippocampus and medial prefrontal cortex (18).

The physiology of memory consolidation can now be investigated using a variety of methods, including pharmacological interventions, closed-loop stimulation, and optogenetics (19-22). When the TMR method was applied during rodent sleep (23), results showed that two sound cues presented during slow-wave sleep differentially biased place-cell firing in accordance with the spatial meaning of those two sounds acquired during prior waking sessions. Although TMR cues influenced hippocampal replay, it was unclear whether changes in memory storage resulted. In humans studied with functional magnetic resonance imaging (fMRI), brain activity during sleep has been elicited by TMR cues, including odors (11, 24) and sounds (25, 26). However, fMRI is not directly sensitive to the primary neurophysiological signals that have been linked with memory processing during sleep, the following three EEG oscillations in particular.

(i) *Slow oscillations* are observable widely across the cerebral cortex, comprising a hyperpolarizing downstate of neuronal silence and a depolarizing upstate corresponding to high levels of neuronal activity. At 0.5-1 Hz, slow oscillations are at the lower end of the slow-wave or delta band, 0.5 to 4 Hz. (ii) *Sleep spindles* are transient thalamocortical oscillations at 12-16 Hz defined as lasting between 0.5 and 2 s. (iii) *Sharp-wave/ripple complexes* include high-frequency ripple activity (70-110 Hz in humans; 27, 28, 29), typically in the hippocampus but also in cortex, and observed contemporaneously with place-cell replay in rodents. These three cardinal oscillations are thought to be temporally coordinated (2). Ripples in the hippocampus tend to be nested in the troughs of sleep spindles, which, in turn, ride on the peaks of slow oscillations (30). Slow-oscillation upstates may provide a synchronized temporal frame that optimizes cross-cortical communication, when specific cortical networks are activated through the guidance of hippocampal-to-cortical communication.

Much remains to be learned about these oscillations in relation to human memory consolidation. Although they do not comprise a complete description of the relevant neurophysiological events, they are currently our best clues about these mechanisms. We also need to understand how the physiology of memory processing during sleep compares with the physiology of memory processing during waking states. Notably, awake EEG signals such as medial temporal theta and gamma oscillations have also been attributed mechanistic roles in memory functions (31).

Manipulating memory processing during sleep with TMR yields the empirical advantage of a precisely timed event that provokes reactivation and concomitant neural activity. Efforts to decipher neural signals of this memory reactivation have shown that it is possible to use EEG signals to implicate specific types of reactivation (32-36). Importantly, to understand how consolidation progresses, memory reactivation during sleep should be liniked with a verifiable improvement in memory storage, usually in the form of improved memory performance after sleep. A key challenge is thus to characterize brain activity during sleep that is causally related to improvements in memory storage.

To examine relationships between human hippocampal activity and consolidation during sleep, we recorded field potentials from electrodes implanted in the medial temporal region (SI Appendix Fig. S1 and Table S1). This rare opportunity to monitor brain activity directly from the hippocampus and adjacent cortex was possible because electrodes were implanted to evaluate neurosurgical treatment options for medically intractable epilepsy. Locations for intracranial EEG monitoring (iEEG) were determined entirely by clinical considerations. Removing the epileptogenic zone with stereotactic laser ablation (37-39) can be very successful in such patients, but it requires localizing the generation of seizures to a specific brain region via iEEG. During this period of monitoring, typically 1-2 weeks in total, patients participated in the following three consecutive phases of the study: (i) an evening learning session, (ii) an overnight period of sleep when TMR sounds were presented, and (iii) memory testing the following morning (see Materials and Methods for procedural details).

We hypothesized that the presentation of sounds during SWS would have behavioral and electrophysiological consequences. Behaviorally, sounds should lead to differential recall for spatial memories associated with those sounds compared to equivalent spatial memories not associated with those sounds. Such memory benefits have been observed using similar behavioral procedures in healthy young individuals (12, 40, 41). Electrophysiologically, we considered this an exploratory study, with the presupposition that hippocampal activity produced in response to learning-related sounds may index the extent of memory change. The sounds presented during SWS included *cues* (sounds from the learning session) and *standards* (a matched set of other sounds that were not part of the learning session for that particular patient, but were for others). Different responses to cues and standards could indicate a difference in memory processing, if the cue sound associated with a specific object would preferentially provoke retrieval of the object’s location. The crucial consequence of this memory reactivation is a selective benefit for remembering the corresponding spatial information after sleep.

## RESULTS

### Memory Performance and TMR Implementation

At the conclusion of the learning procedure, patients completed the pre-sleep test and placed objects reasonably close to the correct locations, thus demonstrating effective spatial learning. The number of learned object locations was adjusted in advance to tax each patient’s abilities at an tolerable level of difficulty (Table 1 shows data for individual patients). The mean spatial error across patients was 193 pixels ±27 (*SEM*). This error corresponds to 4.4 cm from the correct location (roughly the width of each object). Although spatial recall was far from perfect, these errors were much smaller than the level of error expected from random guessing (estimated at 8.9 cm).

**Table 1.** Learning and recall data for each patient.

| Patient | Object locations to learn | Mean runs to criterion<sup>1</sup> | Pre-sleep recall error (cm ± SE) | Sound repeats<sup>2</sup> | Number of sounds | TMR1 & TMR2 trials<sup>3</sup> | Cuing period (min)<sup>4</sup> |
| --- | --- | --- | --- | --- | --- | --- | --- |
| P1 | 20 | 6.9 | 3.1 ± 0.8 | 10 | 200 | 50 | 77 |
| P2 | 10 | 5.4 | 5.6 ± 1.3 | 12 | 120 | 24 | 36 |
| P3 | 10 | 4.4 | 6.1 ± 1.4 | 25 | 250 | 50 | 107 |
| P4 | 14 | 3.9 | 3.9 ± 0.9 | 13 | 182 | 39 | 65 |
| P5 | 10 | 2.7 | 3.3 ± 1.3 | 21 | 210 | 42 | 79 |

*Notes.*
1. Number of runs required to learn each object location, averaged across objects. 2. Number of times each sound was presented during sleep (cue sounds and standard sounds). 3. Number of trials in each category (TMR1, TMR2), following a median split of cues as a function of memory change, omitting the median when there was an odd number of cues. 4. Time from first to last cue presentation, including pauses.

These pre-sleep memory results were utilized in a stratification procedure to select 50% of the objects from learning for TMR in order to match pre-sleep recall accuracy between two conditions. This procedure functioned as intended, in that pre-sleep errors were similar for cued and uncued objects (4.5 cm ±0.7 and 4.2 cm ±0.7, respectively; t_4_=2.05, *p*=0.11).

During sleep, auditory cues were quietly presented for half of the objects from the learning phase and an equal number of similar sounds that had not been used in the learning phase. Sleep physiology from scalp electrodes was monitored online by the experimenter so that sounds would be delivered during SWS and not wake the patient (Table S2 shows offline sleep scoring results). No patients reported hearing sounds during sleep when queried during debriefing. Cue and standard sounds were repeatedly presented with a pause when the sleep stage was no longer SWS or when awakening seemed possible due to extraneous hospital noise. The mean delay from pre-sleep test until TMR was 3 hrs ±1.75 (range 1-5), as it varied with the time required for each patient to fall asleep. The mean delay from pre-sleep test until post-sleep test was 13 hrs ±3.25 (range 10-18.5).

Post-sleep recall showed forgetting, in that patients placed objects 5.1 cm ±1.0 from the correct location, on average, which amounted to a 16% increase from pre-sleep error. As in many prior studies comparing cued versus uncued spatial recall accuracy (13), forgetting was reduced or eliminated for cued objects compared to uncued objects. Fig. 1 shows change in recall accuracy separately for cued and uncued objects. The mean change in error preto post-sleep was -3.19% ±8.8% for cued objects and 32.72% ±10.7% for uncued objects (t_4_=5.97, *p=*0.004, Cohen’s *d*_*AV*_=1.83). Given this relative benefit in recall for locations as a function of whether corresponding object sounds were presented, we next analyzed intracranial EEG data to explore possible relationships with memory processing during sleep.

**Fig. 1.**
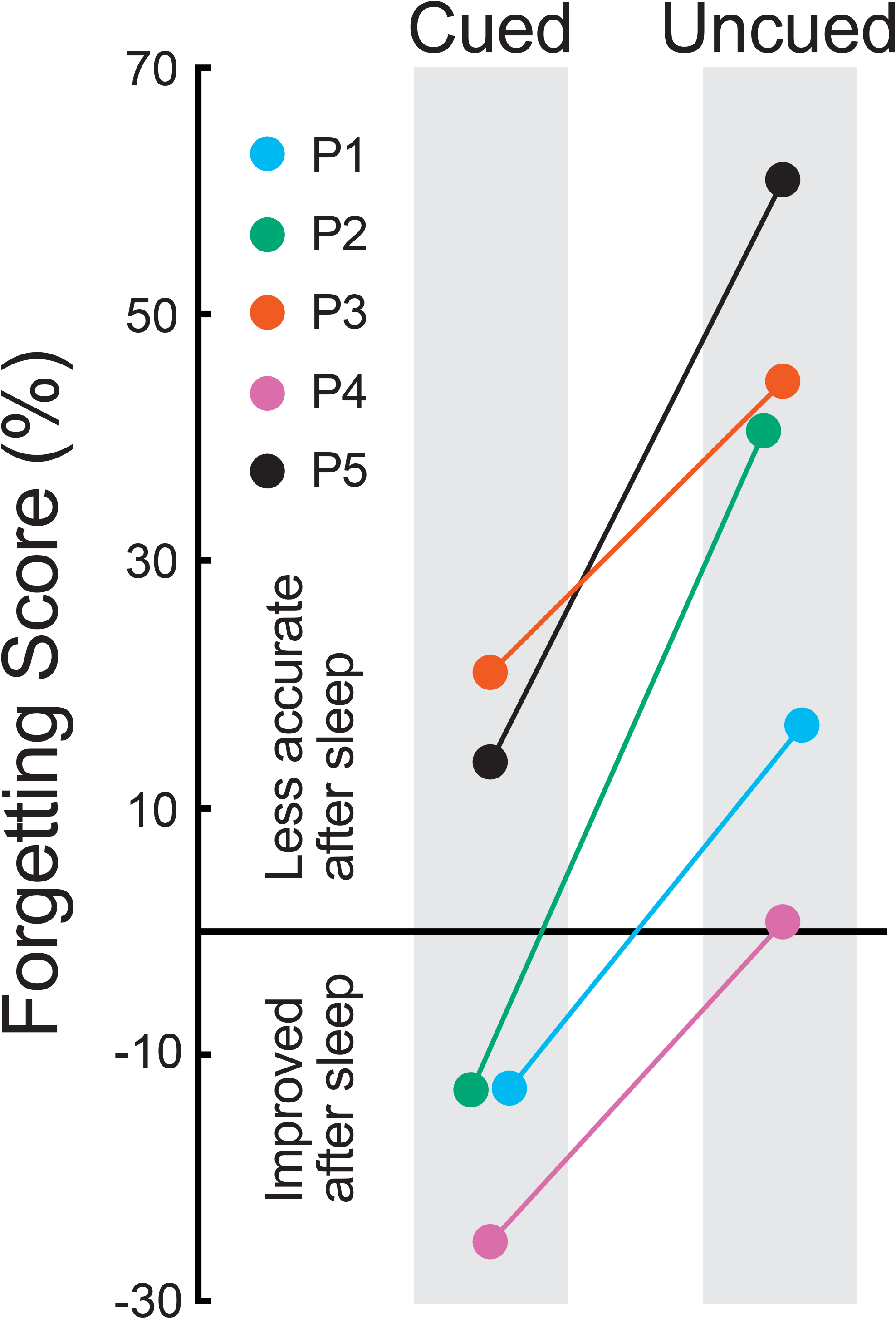
Memory testing results from each patient in the form of forgetting scores, computed as the percent change from the pre-sleep test to the post-sleep test. Positive values reflect reduced accuracy after sleep whereas negative values reflect improvement. Forgetting was observed for uncued objects, as might be expected simply due to the time that elapsed, in that locations were recalled less accurately after sleep. On the other hand, every patient did relatively better for cued objects, and three patients showed an increase in accuracy for cued objects. Thus, the overnight presentation of sounds associated with a subset of the learned objects produced a relative improvement in recall in all five patients.

### Medial Temporal iEEG Responses

Responses to sounds delivered during sleep were analyzed for a broad frequency spectrum based on single-trial event-related oscillatory activity. We interrogated iEEG frequency bands implicated in memory processing in prior studies during wake and sleep: theta (4-8 Hz), sigma (12-16 Hz), and gamma (20-100 Hz). Significant gamma effects were followed up by testing low gamma (gamma_L_, 20-50 Hz), mid-gamma (gamma_M_, 50-80 Hz), and high gamma (gamma_H_, 80-100 Hz, corresponding to ripples). We also conducted separate analyses of ripples, as these discrete events are not necessarily reflected by power in the gamma_H_ band.

Recordings from electrode contacts in the region of the medial temporal lobe (MTL) were examined after first excluding data from contacts within the epileptogenic zone for each patient (as determined via clinical monitoring). To constrain initial analyses, we first identified a single contact for each patient that produced a large time-domain response. For this step, we computed event-related potentials (ERPs) elicited by all sounds presented during sleep (collapsed across cue and standard sounds). We identified one contact for each patient with the maximum ERP peak amplitude between 200-2000 ms after stimulus onset (Fig. S1). Then we identified a cluster (termed *MTL cluster*) of five contacts within or near the hippocampus, surrounding and including the one contact with the largest ERP (Table S3 provides anatomical localization estimates and EEG measures for each MTL contact in each patient, indicating those selected for the MTL cluster). Here, we emphasize frequency-domain responses, which were more consistent across patients than the time-domain responses.

We assessed both how prior learning influenced medial temporal activity during sleep and how this activity influenced memory storage. Primary analyses involved comparing responses to cue sounds versus standard sounds. Sounds in these two conditions (and corresponding objects) were randomly assigned to the two conditions for each patient, so systematic differences in physical stimulus characteristics or associated factors are unlikely to have confounded these comparisons. Each cue sound was associated with a specific object and location during the learning phase. The cue sound was heard repeatedly in conjunction with the object during learning and testing. In contrast, standard sounds had no such connection to recent learning.

### Event-Related Power for Cue Versus Standard Sounds

Time-frequency plots were computed to show event-related power separately for the two types of sounds presented during SWS. If recent memories from the learning phase were not reactivated by cue sounds, then the responses would be expected to be identical for the two conditions. To the contrary, clear differences were apparent for multiple frequency bands, which supports our prediction and the evidence from the behavioral results that cue sounds elicited memory reactivation. The time course of EEG power across the spectrum from 2-120 Hz averaged across trials and patients was produced for both the contact with the largest ERP (Fig. S2) and for the MTL cluster (Fig. S3). A clear divergence in responses between cue sounds versus standard sounds was observed, particularly in lower frequencies. Given that results were very similar for the contact with the largest ERP and for the MTL cluster, we emphasize results from the MTL cluster (Table S3 shows results from all MTL contacts).

An analysis of cue-standard differences for the lower portion of the EEG spectrum is depicted in Fig. 2. Differential responses were apparent at multiple EEG frequencies between 2-20 Hz overlapping with theta and sigma bands. To visualize the time-course of sigma responses, we averaged data over 400-ms intervals (Fig. 2B). For analysis, responses were summed over the post-stimulus interval from 500 ms to 3500 ms, beginning after the stimulus and ending prior to the baseline period of the next stimulus. For each patient, sigma was greater for cues than standards (Fig. 2C). Testing across patients revealed a significant difference between means (cue 0.23 dB ±0.12, standard -0.01 db ±0.13, *t*_4_=6.25, *p*=0.003, Cohen’s *d*_*AV*_=0.93).

**Fig. 2.**
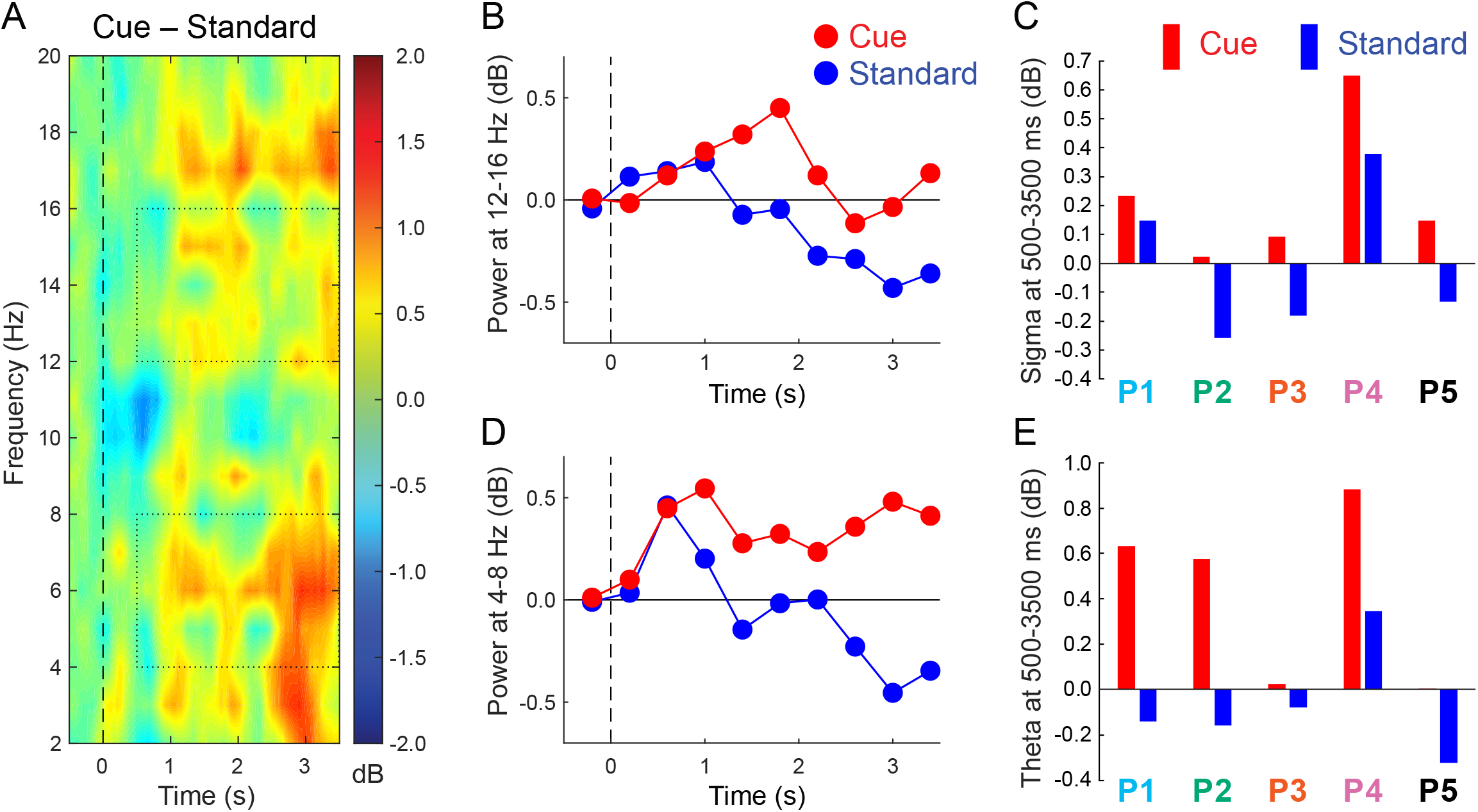
Responses to cue and standard sounds from the MTL cluster. (**A**) Time-frequency plot of the cue-standard differential response across all patients. Dotted rectangles indicate the data used for evaluating responses in the sigma and theta bands, focusing on the interval from 500-3500 ms. (**B**) Sigma measures over consecutive 400-ms intervals, centered over each interval. (**C**) Sigma measures over the poststimulus epoch for individual patients. (**D**) Theta measures over consecutive 400-ms intervals. (**E**) Theta measures over the post-stimulus epoch for individual patients.

A parallel analysis was run for theta. The time-course showed large theta differences between the two conditions over much of the epoch (Fig. 2D). Post-stimulus theta was greater for cues than standards in all five patients, just as for sigma (Fig. 2E), and this cue-standard difference was significant (cue 0.42 db ±0.20, standard -0.07 db ±0.13, *t*_4_=3.94, *p*=0.017, Cohen’s *d*_*AV*_=1.55).

Analyses of cue-standard differences in the gamma band did not reveal significant differences (−0.16 dB ±0.12 and -0.11 dB ±0.09, respectively, *t*_4_=0.49, *p*=0.65). Although differences were apparent in some atients, these effects were not consistent across patients.

To determine whether sigma and theta effects were unique to the MTL, we computed these effects for contacts in other brain regions. We identified a *non-MTL cluster* for each patient, but were limited by the fact that contacts were not placed in the same region in each patient. We thus used other probes or surface strips to select a cluster of 5 contiguous contacts far from the MTL for each patient (located in superior temporal cortex for P1; superior temporal and mid-temporal cortex for P2 and P3; supramarginal cortex for P4; and mid-temporal and lateral occipital cortex for P5). In the non-MTL cluster, the cue-standard contrast was nonsignificant for sigma (cue 0.07 dB ±0.11, standard -0.03 dB ±0.12, *t*_4_=1.57, *p*=0.19) and for theta (cue 0.13 dB ±0.23, standard -0.06 dB ±0.18, *t*_4_=1.11, *p*=0.33). However, the cue-standard contrast did not differ significantly between MTL and non-MTL clusters in either case (sigma *t*_4_=1.75, *p*=0.15; theta *t*_4_=1.55, *p*=0.20).

### Event-Related Power as a Function of Memory Change

Memory change was computed for each object that was cued during sleep by comparing pre-sleep versus post-sleep recall accuracy (forgetting scores, as shown in Fig. 1 across all cued objects). EEG responses were then analyzed for two equal subsets of cues yielding the most benefit from TMR versus the least benefit from TMR (termed *TMR1* and *TMR2* conditions). Mean memory change was -1.63 cm ±0.87 for TMR1 and 1.56 cm ±0.70 for TMR2 (*t*_4_=3.09, *p*=0.037, Cohen’s *d*_*AV*_=1.93). Mean memory error on the pre-sleep test did not differ between conditions (5.68 cm ±1.33 for TMR1 and 4.13 cm ±0.78 for TMR2, *t*_4_=1.05, *p*=0.35). Importantly, memory reactivation presumably occurred and improved memory for both conditions, but with more improvement for the TMR1 condition.

This TMR1-TMR2 contrast, comparing conditions defined by memory change over the sleep interval, revealed differential gamma responses at multiple frequencies (Fig. 3A). Generally, gamma power was greater in association with memory improvement versus forgetting (−0.02 dB ±0.15 and -0.36 dB ±0.18 for TMR1 and TMR2, respectively, *t*_4_=3.87, *p*=0.018, Cohen’s *d*_*AV*_=1.31). These effects can also be visualized in data averaged for the high-gamma and low-gamma band (Fig. 5B-E). For the post-stimulus interval from 500-3500 ms, the TMR1-TMR2 difference was significant at 80-100 Hz (gamma_H_ 0.03 db ±0.12 and -0.29 db ±0.18, *t*_4_=3.00, *p*=0.040, Cohen’s *d*_*AV*_=1.05) and at 20-50 Hz (0.002 dB ±0.18 and - 0.34 dB ±0.17 for TMR1 and TMR2, respectively, *t*_4_=2.97, *p*=0.041, Cohen’s *d*_*AV*_=1.00). Although there were similar trends for mid-gamma, they were inconsistent across patients (gamma_M_ -0.08 dB ±0.15 and - 0.43 dB ±0.23, *t*_4_=2.27, *p*=0.09). Differences were not reliable for sigma and theta bands (sigma 0.36 db ±0.14 and -0.14 db ±0.27, *t*_4_=1.63, *p*=0.18; theta 0.22 dB ±0.23 and 0.49 dB ±0.21, *t*_4_=1.91, *p*=0.13).

**Fig. 3.**
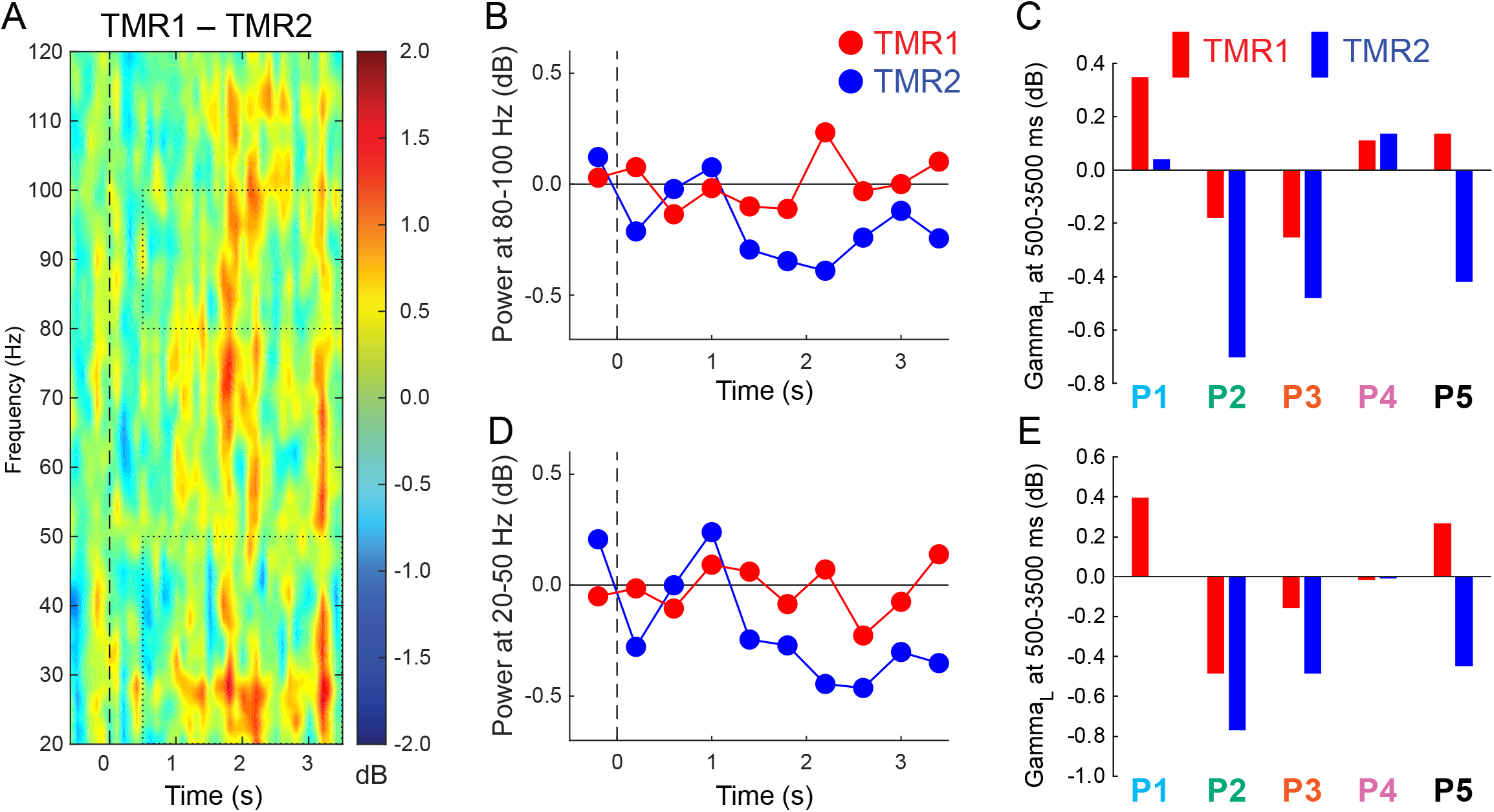
Responses to sounds at 20-120 Hz for two subtypes of the cued condition (larger memory improvement for TMR1 than TMR2). (**A**) Differential responses as a function of memory change (for display purposes, data were smoothed by averaging over 70-ms periods). Dotted rectangles indicate the data used for evaluating responses in two gamma bands, focusing on the interval from 500-3500 ms. (**B**) Gamma_H_ measures over consecutive 400-ms intervals. (**C**) Gamma_H_ measures over the post-stimulus epoch for individual patients. (**D**) Gamma_L_ measures over consecutive intervals. (**E**) Gamma_L_ measures over the post-stimulus epoch for individual patients.

To determine whether these gamma effects were unique to the MTL, we compared TMR1 and TMR2 results for the non-MTL cluster. Differences were nonsignificant for the gamma band (−0.07 dB ±0.10 and -0.26 dB ±0.12, respectively, *t*_4_=1.289, *p*=0.27), although the interaction for MTL versus non-MTL cluster was nonsignificant (*t*_4_=1.02, *p*=0.36).

### Ripples

To expand the frequency-domain analyses in the high-gamma EEG band, we identified ripple activity from the contacts with the largest ERP amplitudes for each patient. We identified a total of 1404 ripples during the cuing period across all patients, and of these, 119 ripples occurred within 3500 ms after a sound onset. For each patient, we compared ripples following cue sounds versus standard sounds on several dimensions. Ripple rate for each patient was the number of ripples identified within 3500 ms of sound onset scaled by number of sounds presented. Ripple latency was measured from sound onset to ripple onset. Ripple duration was determined based on the criteria used to identify ripples. No corresponding cue-standard differences were found (Fig. S4, largest *t*_4_=2.10, *p*=0.10). Likewise, an analysis of TMR1-TMR2 differences also failed to reveal consistent effects across patients (largest *t*_4_=1.22, *p*=0.29 for ripple rate). Relationships between ripple characteristics and memory were also examined using individual cues as the unit of analysis. We determined whether forgetting scores (mean change in error pre-to post-sleep) for each cue sound were correlated with the three ripple parameters, including all patients. No significant correlations were found (all *p*>0.72, strongest *r*(18)=-0.083 for ripple duration).

## Discussion

The primary goal of this research was to gain knowledge about mechanisms of sleep-based reactivation and consolidation of memories through analyses of intracranial recordings. We used the TMR method to provoke memory reactivation, and by reactivating specific memories acquired prior to sleep we were able to analyze iEEG data over time periods when reactivation was likely. Behavioral results showed that comparable memories were less accurate than the selectively reactivated memories, which implies that cues presented during sleep led not only to reactivation of recent spatial memories but also to the sort of memory change generally produced by consolidation.

The memory benefits from TMR shown in Fig. 1 replicate the findings of many prior experiments, some with nearly the same memory paradigm (12, 40), some with small variations in the learning procedure (41-44), and some with related sorts of spatial learning (11, 24, 45, 46). The memory results from these five patients are noteably both for their magnitude and their consistency. Considering the memory benefit from TMR as an advantage in percent change for cued versus uncued memories, patients’ scores were 24%, 26%, 29%, 47%, and 53%, with a mean of 36%. In comparison, this benefit was larger than the 15% advantage found by Rudoy and colleagues (12) and the 13% advantage found by Creery and colleagues (40). In neither of these prior studies did 100% of the participants show the memory advantage for cued over uncued conditions. The learning and TMR procedures were nearly the same, differing primarily in a larger learning requirement (50 object-location associations), testing after a shorter delay (2-3 hrs), and with TMR during an afternoon nap in the quiet environment of a sleep lab. Also, participants were young healthy adults, and their errors in recall were generally smaller than those made by the patients, which may have given patients more room to improve via TMR. In addition, TMR and sleep in general may preferentially benefit weaker memories (40, 47, 48). Patients may also have been more highly motivated to perform well in the learning task because of concern about how epilepsy and its treatment could impact their memory abilities. It is difficult to know which of these factors may be relevant for explaining the larger effects found in the present study. Nevertheless, we can conclude that cues during sleep robustly influenced memory storage in each of the patients, thus setting the stage for a powerful strategy for investigating concurrently recorded brain activity.

Prior studies have linked various aspects of sleep with memory processing, mostly indirectly (4). Some of these studies found correlations between sleep physiology and later memory; others showed that altering brain rhythms during sleep can improve memory upon awakening. In this way, slow waves and spindles were associated with memory improvement. In rodents, hippocampal sharp-wave/ripple complexes and sequential replay of place-cell firing patterns have been linked with memory reactivation during sleep (49, 50). Many of the rodent studies have not provided a direct link with behavioral evidence of memory change. In particular, what sort of memory benefit results from hippocampal replay is unclear, even though we know that hippocampal replay is biased by auditory cues (23). On the other hand, disrupting sharp-wave/ripples had a negative impact on memory (51) and extending them improved memory (52), thus directly linking sharp-wave/ripples with memory change.

The use of TMR here allowed us to ask (a) whether neural activity in the human medial temporal region is associated with reactivation of recently acquired spatial memories and (b) whether this activity varies as a function of memory change. The memory change was a selective improvement for recalling cued object locations, presumably due to better memory for the location and/or the link between the object and its location. We interpreted greater responses to cues compared to standards, and greater responses to TMR1 cues than to TMR2 cues, in a relative manner. Responses were normalized by the pre-stimulus values for each frequency, and many factors can influence the level of activity during the pre-stimulus interval, including factors related to the prior stimulus. Therefore, we make no inferences from the findings that, for some stimulus categories and some patients, responses increased or decreased relative to baseline (Figures 2CE, 3CE). However, baseline factors should not have differed between cues and standards, or between TMR1 and TMR conditions, so these paired comparisons are most informative.

For the first question, we compared responses to cue sounds versus responses to standard sounds. We attribute the differences to memory processing rather than auditory processing per se because both types of sound were randomly selected from the same set of object-related sounds, with only the former presented in the context of the pre-sleep learning procedure. Physical stimulus properties per se are unlikely causes of the differential effects. A reasonable interpretation is thus that the cue-standard effects arose due to prior experiences with the cue sounds during the pre-sleep session. We cannot say whether the cue sounds elicited subjective familiarity or novelty detection during sleep, or if effects were related more closely to implicit or explicit memory. These theta and sigma effects were not sensitive to differences in recall performance among cued objects (TMR1/TMR2), albeit this comparison was made with less statistical power. Nevertheless, we interpret theta and sigma effects as reflections of the different status of cue sounds and standard sounds, and therefore as reflections of differential memory reactivation.

Our primary analyses for each patient focused on activity from the medial temporal region (hippocampus and adjacent cortex). Due to each patient’s unique epilepsy treatment plan, recordings were available from different brain locations in each patient, but all patients had at least one medial temporal probe and data from locations determined to be epileptogenic were omitted. We used the independent method of measuring maximal ERP amplitudes for each patient to select a medial temporal contact combined with adjacent medial temporal contacts to constitute the MTL cluster for primary analyses.

Localizing contacts anatomically is one aspect of understanding these data, but another challenge is to determine the sources of the electrical signals in question, given that potentials volume-conduct through the brain. One estimate is that contacts are sensitive primarily to potentials generated within 2 mm (39), but in some circumstances activity may travel further. In studies of P3-like potentials, for example, activity taken to be generated in the hippocampus was recorded in hippocampal contacts as well as in nearby non-hippocampal contacts (53). Here, the consistency of the theta, sigma, and gamma effects is suggestive of similar neurophysiological processing across patients, despite the divergent locations of contacts. Note that regions of entorhinal and parahippocampal cortex are likely relevant for consolidation through their hippocampal connectivity (16, 54), and contacts in these regions may also be sensitive to fields produced in the hippocampus. In this study, anatomical precision is limited by the sparse sampling of electrical activity provided by depth probes, the large size of the probe contacts (1.25 × 2.5 mm), and the possibility that multiple medial temporal regions generate activity in association with consolidation. Further investigation with microelectrodes that can record from specific layers in these brain regions, in patients or in nonhuman animals, would be needed to support stronger conclusions about the neuroanatomical sources of the iEEG effects we described.

We can nevertheless state that the medial temporal responses demonstrated systematic differences for cue sounds compared to standard sounds. Larger responses were found in multiple bands in the seconds following sound onset. Given the literature on oscillations in the theta and sigma bands in relation to medial temporal memory processing, it is striking that these two bands both showed robust effects. The cue-standard effects for both theta and sigma were found in each of the five patients.

Theta in the human temporal lobe, for example, was found to increase just prior to word recall, along with high-frequency activity (55), and several studies have established a link between TMR and theta in human scalp EEG (56-58). One view about theta is that it regulates hippocampal information flow by entraining gamma (59). Intracranial cortical theta has been observed prior to hippocampal SWRs in humans, interpreted as evidence that theta plays a special role in coordinating hippocampal-neocortical dialogue (60). The broader literature on theta and memory, including both scalp and intracranial EEG studies, has mixed findings, and theta increases have not always been associated with memory improvement (61). In our study, theta showed a preferential increase following cue sounds, presumably related to memory reactivation, but we did not find a relationship with magnitude of TMR benefit. A speculative interpretation consistent with the literature is thus that the medial temporal theta increases we observed function to coordinate the strengthening of memories, which itself is more directly reflected by gamma activity (as discussed below).

Sleep spindles, which have been linked with memory function (62), intersect with the sigma band investigated here (12-16 Hz). Two types of spindles have been distinguished, slow spindles at 11-13.5 Hz and fast spindles at 13.5-16 Hz (63). In a previous study with the same TMR procedures, we found that fast spindles after cues predicted a relatively better spatial memory improvement (40). Similar spindle relationships have been observed in other TMR studies (42, 64). Interestingly, restoration of sleep spindles with the treatment of a childhood epilepsy syndrome has been correlated with improvement in cognition and memory (65). Spindles are classically defined in scalp EEG, so it remains unclear how they might be related to the medial temporal sigma effects here. We speculate that the sigma effects could have occurred concurrently with spindles in various cortical regions in association with memory reactivation, but our ability to observe concurrent cortical spindles was very limited. Additional insights could be gained by considering spindles and slow oscillations, particularly given that spindles synchronized with slow-oscillation upstates appear to facilitate memory consolidation (32, 64, 66). For example, TMR sounds could be presented at specific times to engage spindle processing (64), or relatedly, concurrent with slow-wave up-states (22, 67, 68).

Together, the theta and sigma effects can be attributed to additional processing provoked by sounds due to the learned associations. This additional processing likely pertains to the multiple pre-sleep episodes when each sound accompanied to-be-memorized object locations. Because of the improvement in spatial recall that resulted from TMR, it follows that cue sounds during sleep likely provoked retrieval of object-location associations. Although there may have been no subjective experience of retrieval, this retrieval can be likened to visual imagery for the object in its proper place on the computer screen during learning. We thus attribute the theta and sigma increases to spatial memory reactivation.

Given this evidence of memory-related increases in sigma and theta, we also looked for electrophysiological correlates of memory change. We took advantage of the recall results showing that TMR produced memory improvement for cued objects and that the improvement was not the same for every object. Through a median split, we created two conditions, TMR1 and TMR2, that differed in the degree to which recall for cued object locations changed from pre-sleep to post-sleep. TMR1 memories improved the most overnight; TMR2 memories showed less improvement. In both conditions, memory was better after sleep compared to the uncued condition. The TMR1/TMR2 contrast allowed us to determine whether brain activity systematically differed in association with memory change. Indeed, power in the gamma band after a cue was greater with the superior memory benefit.

We focused analyses on multiple portions of the gamma band, including the frequency of ripple events. Prior studies analyzed gamma oscillations in the range from 25-140 Hz (31), and gamma in general is thought to represent local processing in the cortex (69). Ripples have been found in the hippocampus, adjacent cortex, and other parts of the cortex, mostly in rodents but in a few recent human intracranial studies (27, 28, 30, 70). Hippocampal ripples probably precede medial temporal cortex ripples, in keeping with hippocampal output reaching adjacent cortex next on its way to other cortical regions (27, 71, 72).

Our analysis of ripples failed to reveal an association between ripples and either differential cue-standard processing or differential processing based on memory change. Despite the significant gamma_H_ effect for TMR1-TMR2, ripple results were not consistent across patients, although there was also variability in contact location across patients and in some patients the ripple analysis was not conducted for a hippocampal contact. Whereas P1 showed a larger gamma_H_ response for TMR1 compared to TMR2, hippocampal ripples in this patient did not show an increase for TMR1 compared to TMR2. Yet, these null findings for ripples do not support strong conclusions, given that associations between ripples and memory might be evident if a larger number of sounds were presented or a larger group of patients tested.

The similarity across patients in three critical bands — increased theta in the cue-standard contrast, increased sigma in the cue-standard contrast, and increased gamma in the TMR1-TMR2 contrast — in conjunction with the consistent memory benefits of TMR across patients argues that the iEEG results may reflect important memory processing. Our interpretation is that these results reflect processing of memories of pre-sleep spatial-learning episodes in the hippocampus and medial temporal cortex. Although these effects were not found at all locations in the brain, studies with broader coverage and additional methods are needed to more fully specify the contributions that many brain regions may play in memory processing during sleep.

Additional limitations that may restrict the generalizabily of the present results include the type and duration of the patients’ seizure disorders, their cognitive capabilities, and other personal characteristics. Whereas a larger sample would have been preferable, that was not possible for this study. In addition, patients with medically intractable epilepsy have a history of anti-seizure medications plus potential seizure-related reorganization of networks (73) and many have neuropsychological deficits. Patients with hippocampal damage, as is found in seizure-related reorganization, have reduced slow-wave activity at the scalp, but slow and fast spindles are indistinguishable from control patients (74). Whereas the consistency of results across the five patients suggests that the observations were not artifacts of a low sample size, replication in additional patients and convergence across multiple methods would be useful.

To summarize, the present intracranial recordings provide a bridge to many other types of neuroscientific research that can advance our understanding of memory consolidation. By combining iEEG with the TMR method, neural activity in multiple frequency bands was linked with memory processing during sleep.

Theta and sigma activity were engaged in conjunction with memory reactivation, which precipitated improvements in memory storage, as demonstrated by clear post-sleep improvements in spatial recall. The results further linked gamma activity with the magnitide of these memory benefits. We thus propose that activity in the lower frequencies reflected recapitulation of object-related memories, allowing for coordinated processing across brain regions, whereas the higher-frequency effects (gamma) reflected improved storage of the associated spatial information. Overall, the findings contribute to our understanding of neurophysiological mechanisms of memory by elucidating some of the neurocognitive steps whereby storage is improved during sleep.

## Materials and Methods

### Participants

Patients underwent invasive EEG monitoring at the University of Chicago Epilepsy Surgery Unit as part of a clinical epilepsy surgery evaluation. Nine patients were tested with the experimental procedures described below over the course of one evening and overnight sleep, concluding with memory testing in the morning. We report results from five of these patients, focusing on responses in the medial temporal region. We excluded results from four other patients because focal medial temporal seizure activity overnight precluded analysis of responses to sound cues from these brain locations. This study was approved by the Institutional Review Boards of the University of Chicago and Northwestern University, and patients gave informed consent. Demographic and electrode implantation information is shown in Table S1.

Following surgical implantation of recording electrodes, electrical activity and video were continuously monitored to observe the onset and progression of seizure activity. Depth electrodes targeted the hippocampus on one or both sides (Integra LifeSciences, Princeton NJ) and surface electrodes were placed over regions of the cortex selected individually for each patient. Electrodes were also placed on the scalp at 24 locations (Cz, C3, C4, Fp1, Fp2, Fz, F3, F4, F7, F8, F9, F10, Pz, P3, P4, P7, P8, T7, T8, T9, T10, Oz, O1, and O2). Recordings from scalp and intracranial electrodes were amplified and acquired with a low-pass filter at 344 Hz and a sampling rate of 1024 Hz.

### Procedure

The experiment began several days after electrode implantation, when patients were willing to participate. Memory testing was conducted using a laptop computer in the patient’s hospital room. The experimenter gave verbal instructions to the patient, and then walked through a shortened version of the task to assure that the procedure was understood. The experiment included four phases: Learning, Pre-Sleep Test, Sleep with TMR, and Post-Sleep Test.

The patient was asked to learn the locations of a set of common objects shown on the computer screen (Table S4). Each object was shown with a concurrent sound closely associated with the object (e.g., picture of cat shown with meow sound). For each patient, 10-20 object-sound pairs were randomly selected from a set of 30 pairs from a prior TMR experiment (40). Half as many additional objects were randomly selected from the set for each patient so that the corresponding sounds could be used as standard sounds during sleep. Table 1 includes additional details for each patient.

### Learning

The learning phase began 90 min before the patient’s expected bedtime. First, the patient was asked to view the objects and try to memorize each of their locations (randomly selected for each patient). Objects were shown on the screen superimposed on a background grid, a 10 × 8 array of squares colored red, blue, or grey, subtending 1144 × 704 pixels (26 × 19.5 cm). Each object appeared on a square white background (4.5 × 3.4 cm) with a red dot superimposed at the center. Objects appeared for 3500 ms, separated by a 1000-ms interstimulus interval. Object onset was always synchronized with corresponding sound onset. Sounds were 500 ms in duration.

After viewing each object once, the patient was asked to recall the object locations. Each trial began with an object appearing in the center while its corresponding sound played. The patient re-positioned the object using a computer mouse, clicking the mouse when the object was in the location they recalled. Feedback was provided at that point, as the object jumped to the correct location and the corresponding sound played again. After 3500 ms, the next recall trial began. Patients were progressively able to place objects more accurately, learning from the feedback.

Learning continued with the following drop-out scheme. The objects were presented in a series of runs, using a different random order for each run. When an object was placed within 3.4 cm of the correct location (i.e., the height of each object), the response was considered correct. If an object was placed correctly in two consecutive runs, it did not appear again during the learning phase. The remaining objects were shown until the drop-out criterion was satisfied for all objects. Achieving this criterion required an average of 4.7 runs.

### Pre-Sleep Test

After an approximately 30-min break, there was an assessment of location recall before sleep. Recall was tested using the same format as in the learning phase, except each item was tested only once and no feedback was given. Individual objects were ranked by recall accuracy so that two sets of objects could be selected via an automated stratification algorithm. That is, objects were assigned to cued and uncued sets in order to closely match the two sets for mean recall accuracy on the pre-sleep test.

### Sleep with TMR

When the patient was ready for sleep, white noise was presented from a speaker located 1 - 3 feet from the patient’s head. Given that background noise in the hospital was louder and less predictable than in the sleep lab where we ran prior TMR experiments, sound intensity was adjusted here on an individual basis to be as low as possible while still being audible above the background. The experimenter remained in the hospital room to monitor sleep physiology from scalp electrodes until TMR was completed. When the experimenter determined that SWS had been reached for at least two 30-s epochs, sound presentation was initiated. Selected cue sounds were randomly intermixed with an equal number of standard sounds and played through the speaker at the same intensity as the white noise.

Sounds presentation was halted if a shift of sleep stage was observed, such as when slow-wave amplitude decreased, or if there was an interruption due to clinical monitoring or other activity. For each set of sounds, one sound played every 4.5 s until all sounds had been presented (with the exception of P1, when the interval was jittered between 3.8-12.5 s). After collecting data from P1, we adopted a constant interstimulus interval as in prior TMR studies (12,40,41). A short pause followed the presentation of all the sounds (mean 3 min), and then if SWS continued based on online inspection, the sounds were repeated in a new random order. Offline sleep scoring confirmed that 96.4% of the sounds were presented during SWS and virtually all of the others during N2 sleep. Table 1 shows the total number of sounds presented for each patient.

### Post-Sleep Test

The next morning, memory was tested for the object locations. As in the Pre-Sleep Test, each object appeared in the middle and the patient re-positioned it to the remembered location. Following the test, patients were debriefed to determine whether they thought they heard any sounds overnight.

## Data Analysis

### Behavior

Spatial recall accuracy was compared between cued and uncued objects to determine effects of TMR. Recall error was computed as the Euclidean distance between each object’s correct location and where it was placed on each test. Forgetting scores were computed as the mean percent change from pre-sleep to post-sleep. The TMR effect was computed as the difference in forgetting score for cued and uncued objects. A median split of cued-object forgetting scores yielded two subcategories, TMR1 and TMR2 (objects with the most benefit from TMR vs. the least benefit from TMR, respectively). Analyses were conducted using paired-sample *t*-tests (two-tailed, alpha = 0.05).

### MRI and CT

To localize electrodes, we used a preoperative T1-weighted MRI and a postoperative CT scan. CT scans were registered to MRI scans using the mutual-information method through the Statistical Parametric Mapping toolbox in MATLAB (75), and cortical reconstruction and volumetric segmentation was performed with Freesurfer (76). Electrode contacts were localized using in-house software (77), which segments electrodes from the CT based on intensity values. The anatomical location of each contact was identified based on the nearest label in the aparc parcellation, generated for each patient through Freesurfer.

### Scalp EEG

Scalp recordings were re-referenced to the common average from all scalp electrodes for each patient. For online sleep staging, data were filtered at 0.1 - 70 Hz with a 60-Hz notch filter. EEG preprocessing was carried out with MATLAB and EEGlab (78). Offline sleep scoring was completed with sleepSMG (http://sleepsmg.sourceforge.net) blind to when sounds were presented.

### Intracranial EEG

Recordings from electrodes in the presumed seizure onset zone were excluded from all analyses (identified based on abnormal rhythmic activity during focal seizures or the earliest activity during the transition from focal to bilateral tonic-clonic seizures). We also removed noisy channels with variability greater than 5 SD compared with all electrodes, and trials with variability greater than 3 SD compared to all other trials. Data were then re-referenced to the common average for each participant. Data were high-pass filtered at 0.01 Hz and notch filtered at 60 Hz and its harmonics to remove slow drift artifacts and line-noise, respectively.

For medial-temporal-lobe depth-electrode recordings during sleep, time-frequency decomposition was conducted using Morlet wavelets. Data were epoched from 1000 ms before each sound to 3500 ms after the onset of each sound. Baseline values, defined as the average spectral power from -500 to 0 ms, were subtracted separately for each frequency. The stimulus-locked time series was normalized using a decibel conversion. Event-related spectral responses were measured at canonical frequencies using Morlet wavelets, computed from 2-120 Hz in single linearly spaced steps, including from 3-20 cycles also increasing linearly as frequency increased. To visualize the full spectrum, wavelets were computed similarly except using 60 logarithmically spaced steps. For visualization of the time-course, data were averaged across 400-ms intervals beginning 400 ms prior to stimulus onset, placing each value at the center of the interval (e.g., -200 ms, 200 ms, 600 ms). To cover the majority of the epoch after the stimulus and avoid edge artifacts, analyses of theta, sigma, and gamma frequencies used mean power averaged over the 500-3500 ms interval.

### Ripples

Ripple identification was conducted using the following steps (30). Data were filtered at 80-100 Hz. Root-Mean Square (RMS) values were computed for 20-ms intervals sliding by 1 ms. A putative ripple was identified when RMS exceeded the 99^th^ percentile for RMS for 38 ms or more (three 80-Hz cycles). If two ripples were separated by <10 ms, they were combined. The final requirement was using MATLAB findpeaks on unfiltered data (smoothed by averaging each 3 samples) to observe at least three peaks and three troughs during the putative ripple period. Data from trials with artifacts from interictal spikes or high-frequency noise were removed from the analysis.

### Data Sharing Plan

Datasets and custom code supporting the current study will be deposited and released on project completion (https:/deepblue.lib.umich.edu/data/).

## Acknowledgements

We thank Susan Florczak, Bruce Caughran, Mallika Patel, and Satoru Suzuki for valuable assistance with many aspects of this research. We are also grateful to the patients and their families who so generously gave their time and effort. This research was supported by the National Science Foundation (BCS-1829414 and BCS-2048681) and the National Institutes of Health (T32NS047987 and T32HL007909).

## Supplementary Information for

**Fig. S1.**
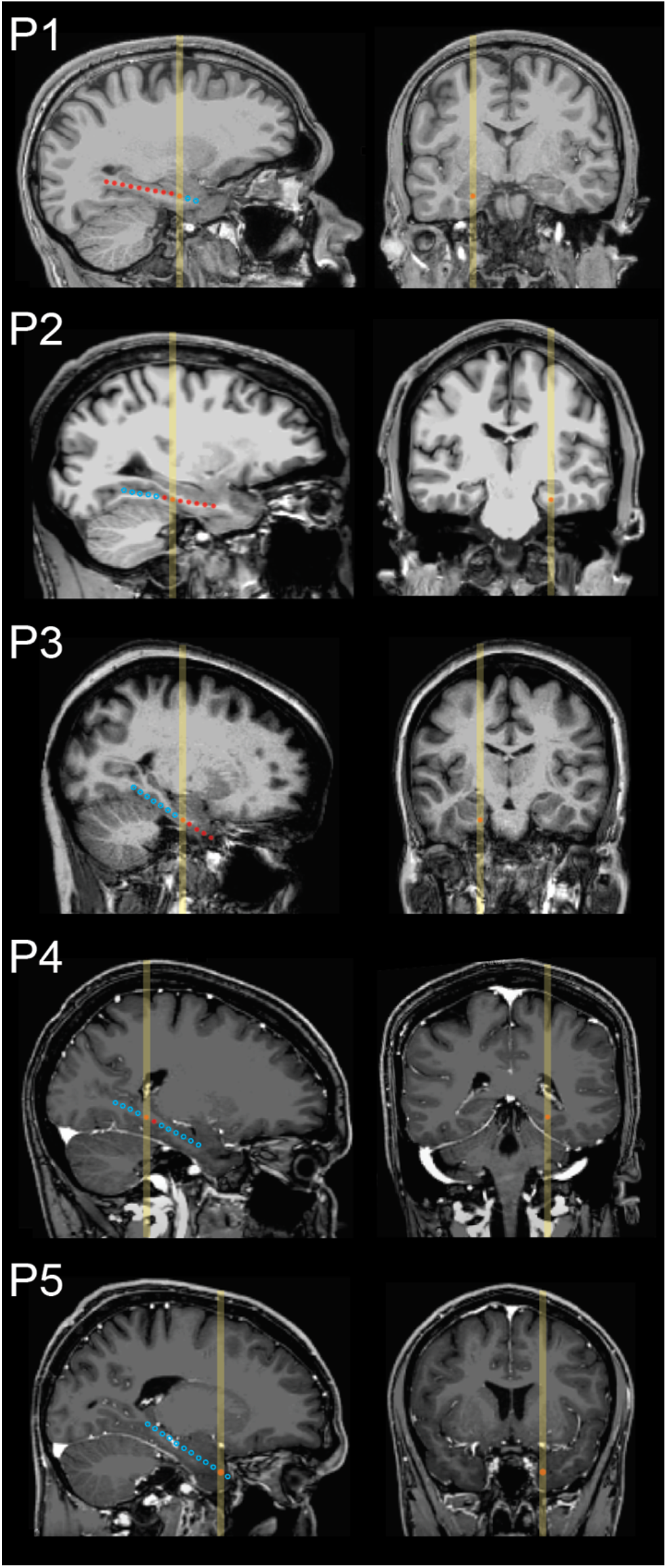
Locations of intracranial electrodes used to monitor medial temporal activity in patients P1-P5. Depth probes were implanted through a small hole in the skull and targeted for the hippocampus, guided by anatomical information from pre-surgical MRI. Contacts were cylindrical, 1.25 mm in diameter and 2.5 mm long, situated 5 mm apart along each probe (measured center-to-center). Images on the left show a sagittal section that includes the estimated location of the contact selected for primary iEEG analysis as an orange circle. Locations of other contacts within the section are shown as red circles. Locations of contacts in adjacent sagittal sections were projected onto approximate locations in the section and shown as open blue circles. The yellow vertical line on each sagittal and coronal image indicates the location of the selected contact and corresponds to the approximate level of the section opposite.

**Fig. S2.**
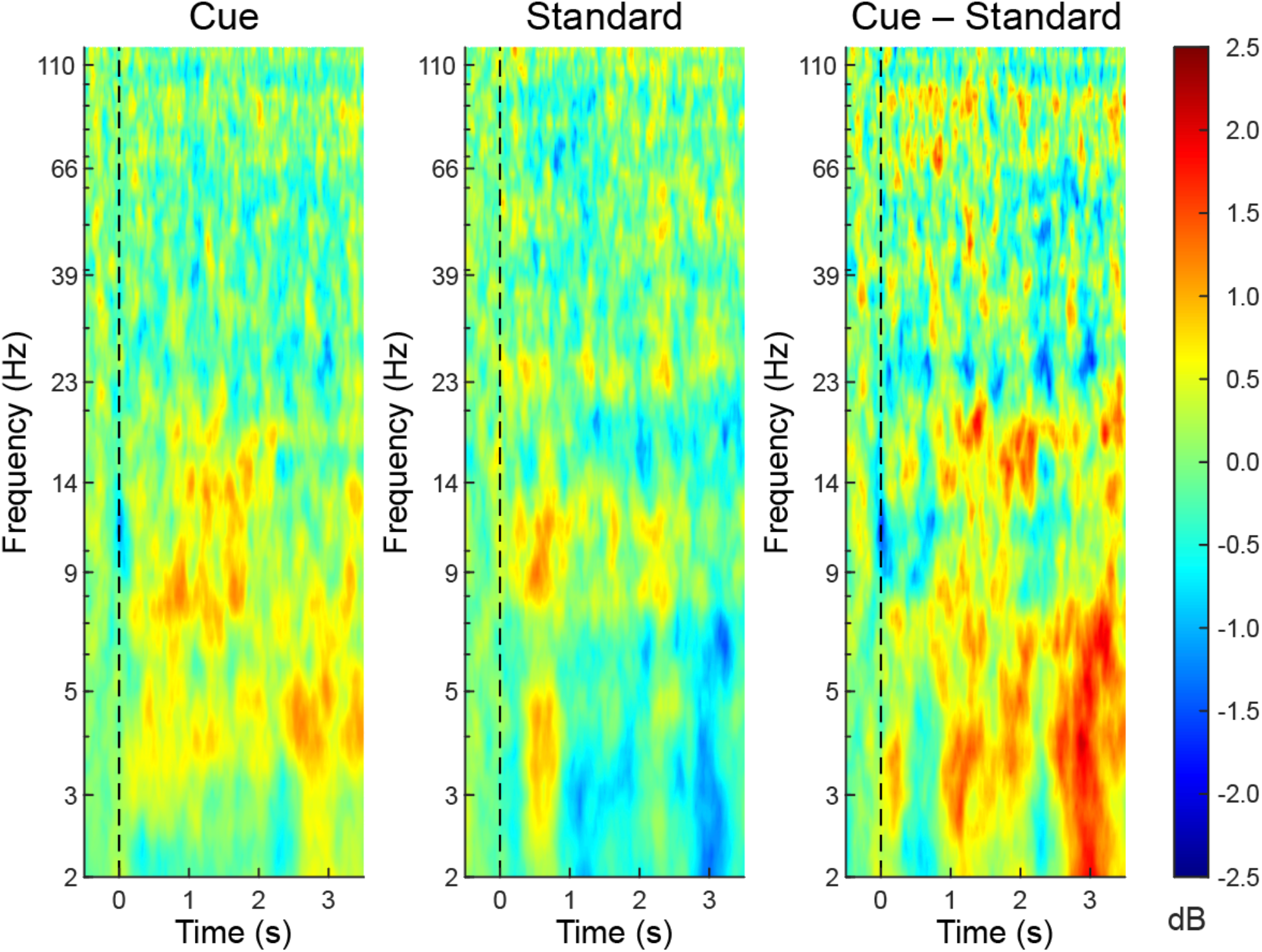
Responses to sounds during sleep from the medial temporal location with the largest ERP amplitudes in each patient (locations and ERP values listed in Table S3). Time of sound onset indicated by dashed vertical lines (0 ms). Intracranial EEG responses to each sound presentation were analyzed to yield time-frequency responses across frequencies from 2-120 Hz (log scale), averaged for each condition within each patient, and then averaged across patients. Color indicates dB power, baseline-corrected using mean power over 500-ms interval prior to sound onset. Responses differed according to whether sounds were those used during the spatial learning task (cue sounds, left panel) or sounds not used during the spatial learning task (standard sounds, middle panel). Differences can be seen in the subtraction of responses to standard sounds from responses to cue sounds (right panel).

**Fig. S3.**
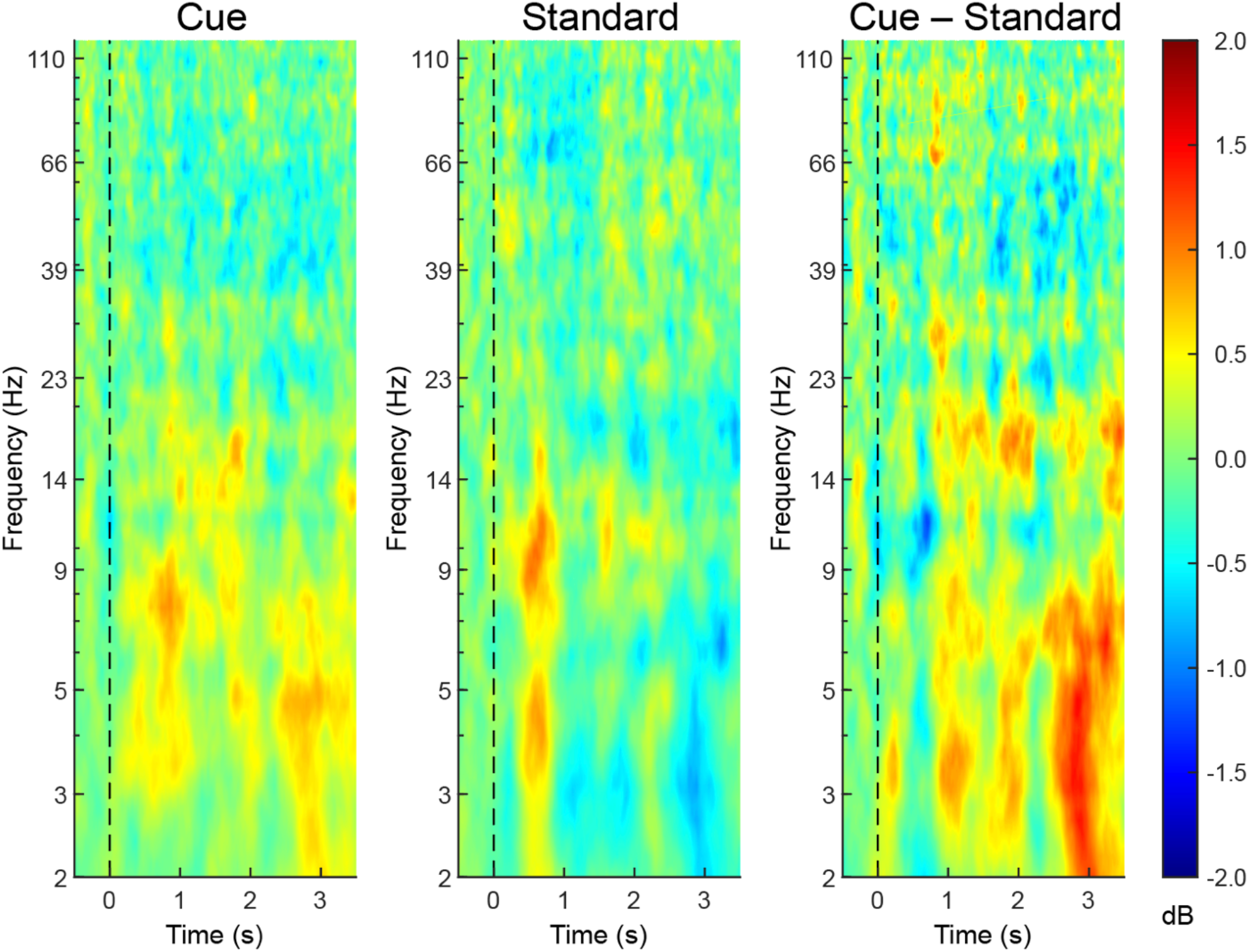
Responses to sounds during sleep averaged over a set of five medial temporal locations (see Table S3), following the same format as in figure S2. Time of sound onset indicated by dashed vertical lines (0 ms). Intracranial EEG responses to each sound presentation were analyzed to yield timefrequency responses across frequencies from 2-120 Hz (log scale), averaged for each condition within each patient, and then averaged across patients. Color indicates dB power, baseline-corrected using mean power over 500-ms interval prior to sound onset. Responses differed according to whether sounds were those used during the spatial learning task (cue sounds, left panel) or sounds not used during the spatial learning task (standard sounds, middle panel). Differences can be seen in the subtraction of responses to standard sounds from responses to cue sounds (right panel).

**Fig. S4.**
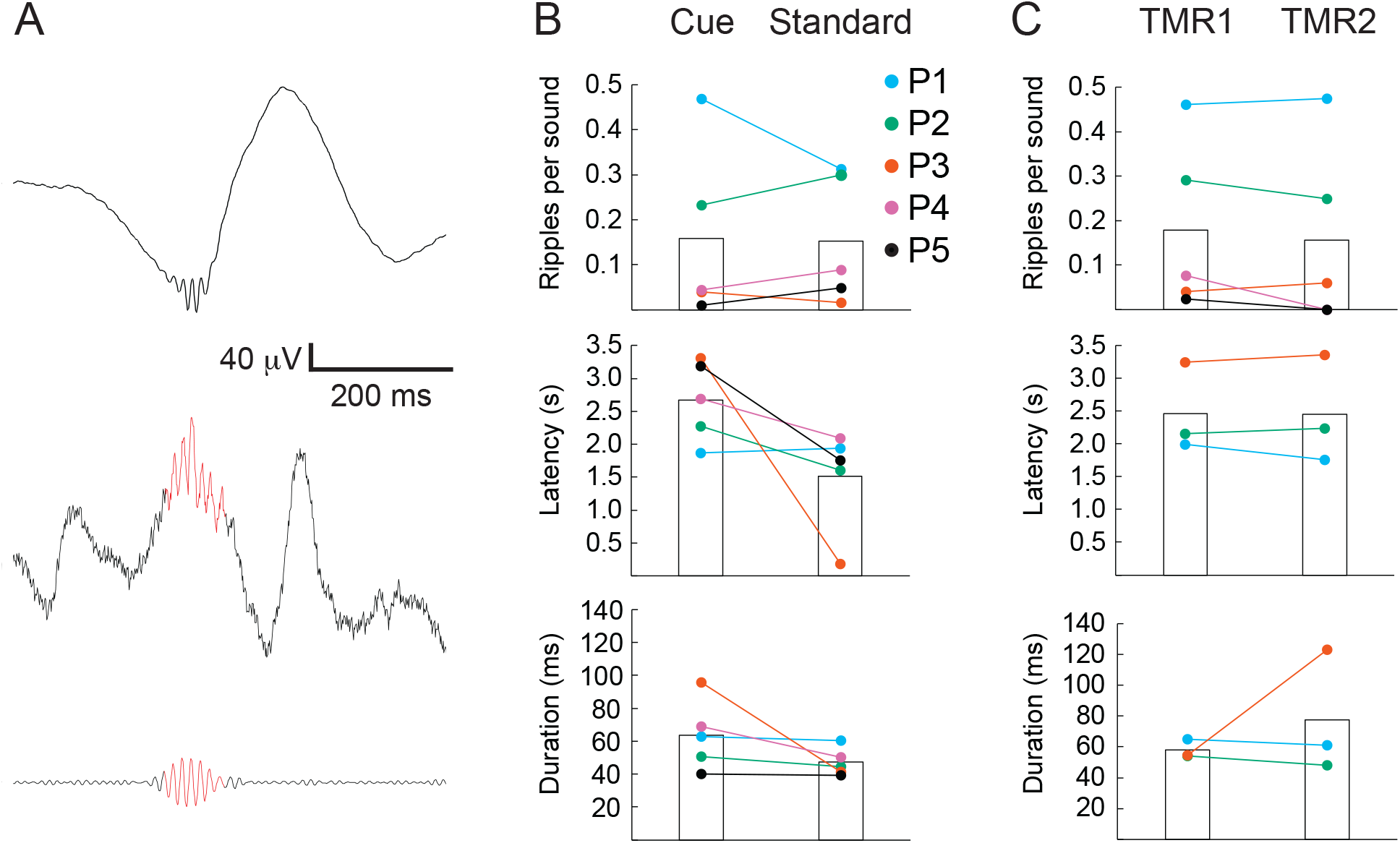
Ripple results obtained from the one contact in each patient with the highest ERP amplitude to all stimuli during sleep. (A) As a representative example, in one patient (P1) a total of 396 ripples were identified, averaged time-locked to the largest positive peak, and then high-pass filtered at 3 Hz (top). A single ripple from the same patient is shown below (60-Hz notch filter, ripple in red) along with the same data filtered at 80-100 Hz (bottom). (B) Comparisons for ripples elicited by cue versus standard sounds during sleep showed no consistent effect on ripple rate, latency, or duration. (C) Comparisons between trials separated as a function of change in memory from before to after sleep also showed no consistent differences in ripple rate, latency, or duration. Because two patients showed no ripples for one condition, their data were not included in the latency and duration analyses.

**Table S1.**
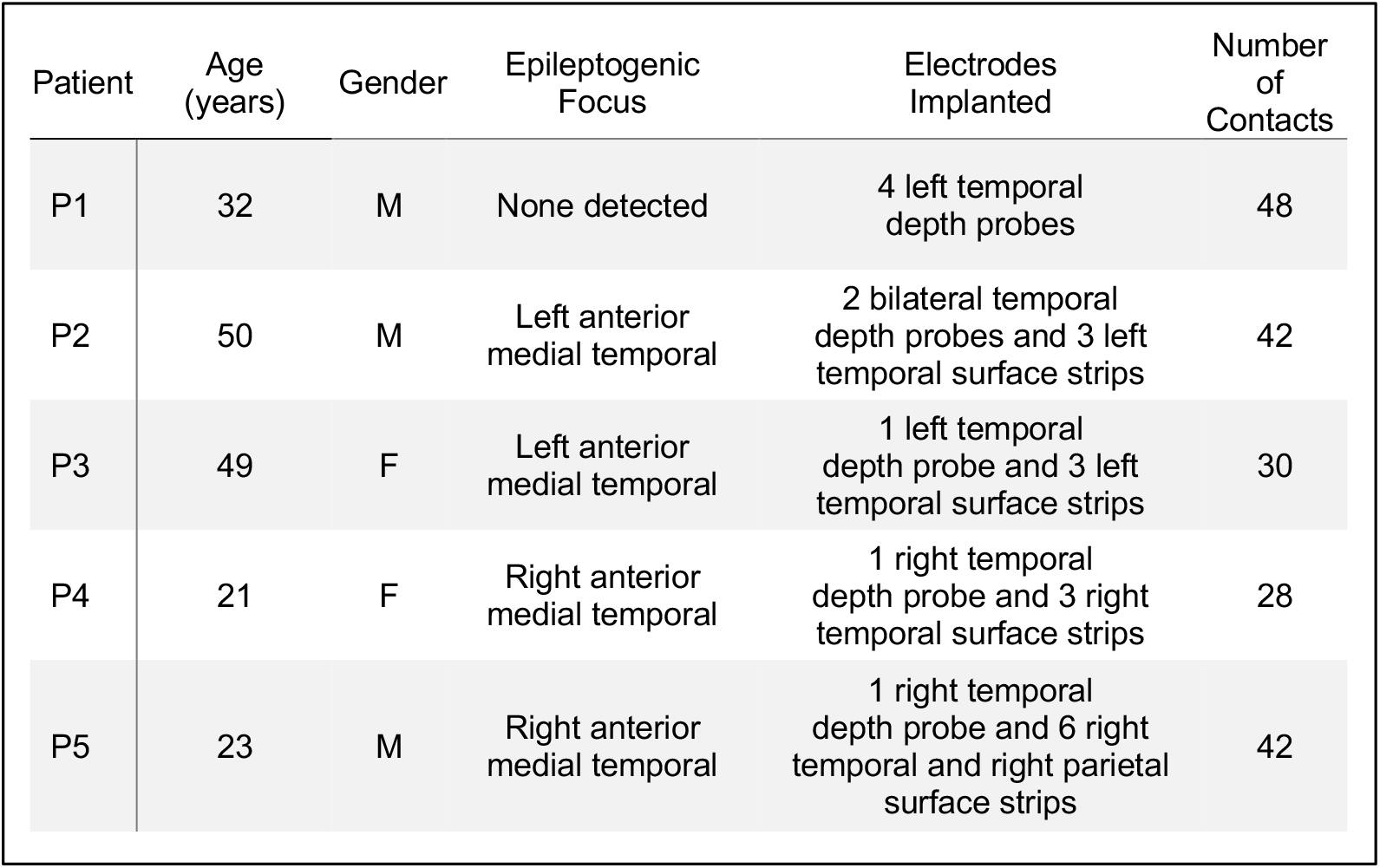
Individual patient demographics and electrophysiological monitoring details.

**Table S2.** Sleep summary across patients based on scalp EEG data (values were based on best estimates for sleep staging, given that recordings were not typical polysomnographic recordings, sometimes with low-quality EEG and lacking electro-oculographic and electromyographic channels).

| Stage | Mean (min) | SE (min) | Range (min) |
| --- | --- | --- | --- |
| N1 | 46.7 | 8.2 | 24.5 – 61.5 |
| N2 | 158.2 | 23.0 | 85.5 – 227.5 |
| N3 | 93.5 | 14.8 | 52.5 – 137.5 |
| REM | 28.2 | 5.3 | 10.5 – 40.0 |
| Total | 326.6 | 27.7 | 222.0 – 385.5 |

**Table S3.**
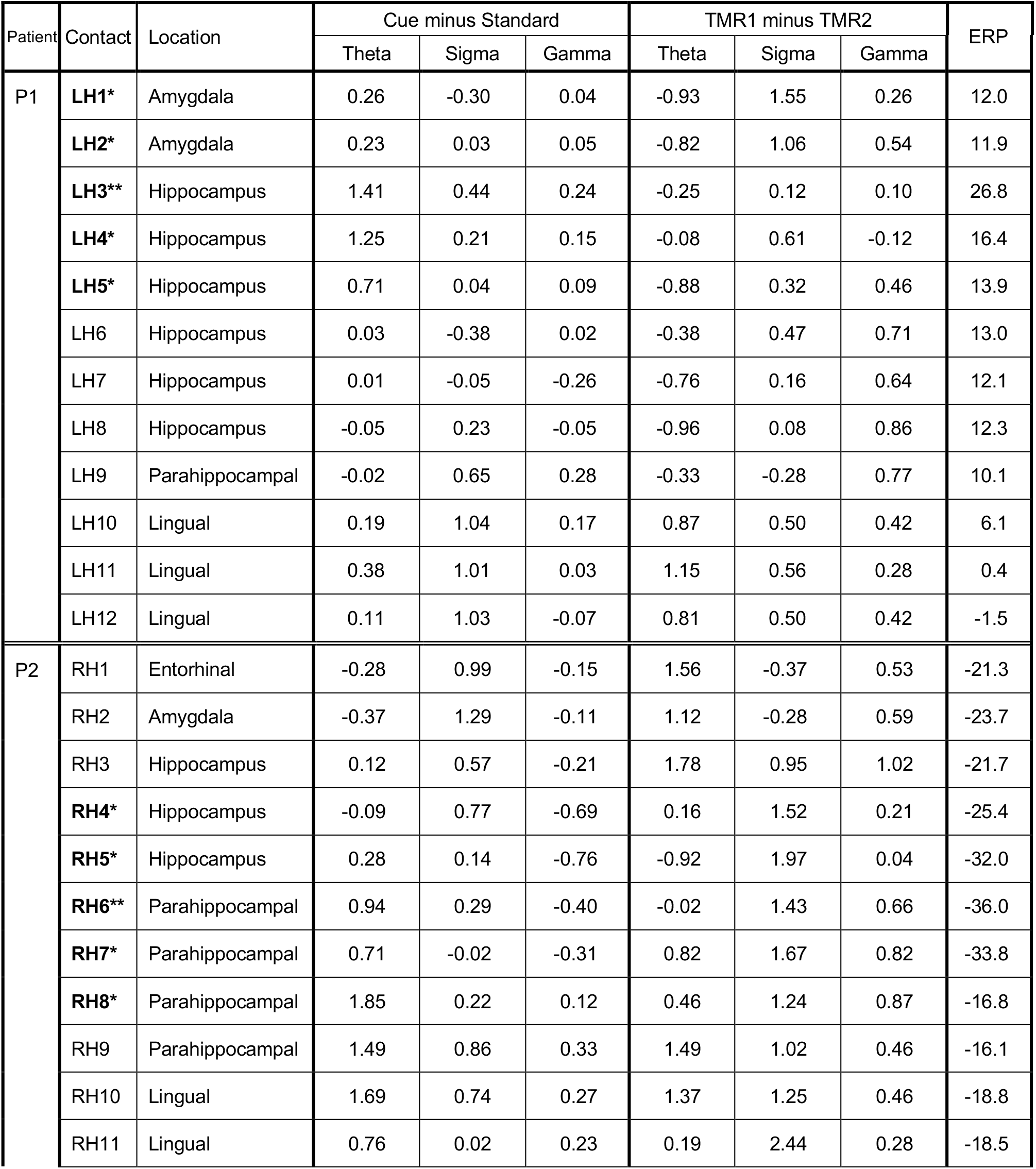
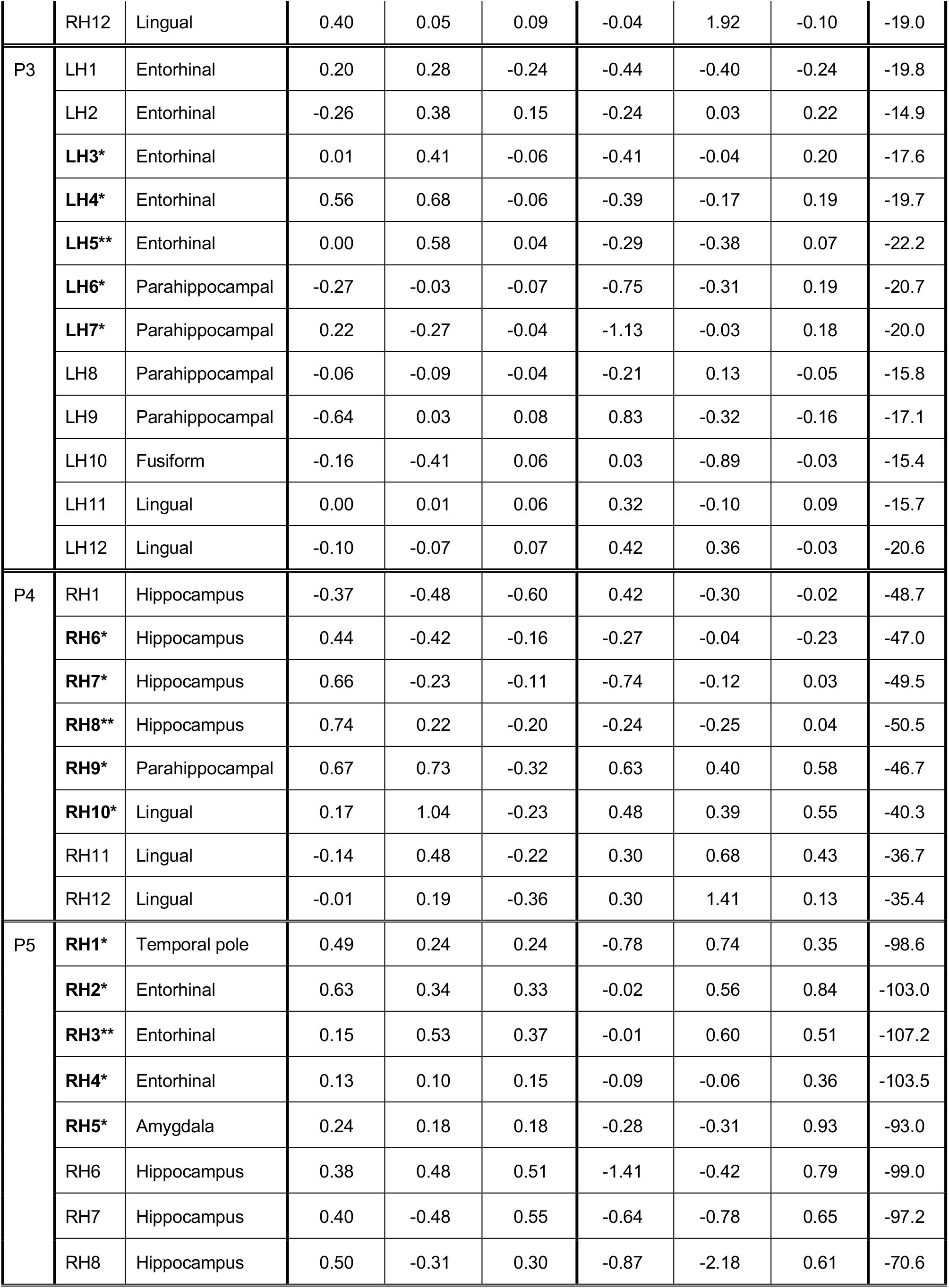
Data from medial temporal probes for each patient (P1-P5). Contacts were on probes either in the left hemisphere (LH) or in the right hemisphere (RH). The five contacts used for the MLT cluster are shown in bold (with ** for the contact with the largest ERP amplitude and * for the other four contacts). Contacts RH2-5 in patient P4 and RH9-12 in patient P5 were omitted from analysis due to excessive seizure activity. Each contact was located within or close to the cortical region or brain structure listed. Measures are shown for differential EEG power in dB, baseline-corrected, for theta power (4-8 Hz), sigma power (12-16 Hz), and gamma power (20-100 Hz). Lastly, ERP amplitudes (in microvolts) are shown for the maximal baseline-to-peak amplitude between 200-1200 ms after stimulus onset (collapsed for cue and standard stimuli delivered during sleep).

**Table S4.**
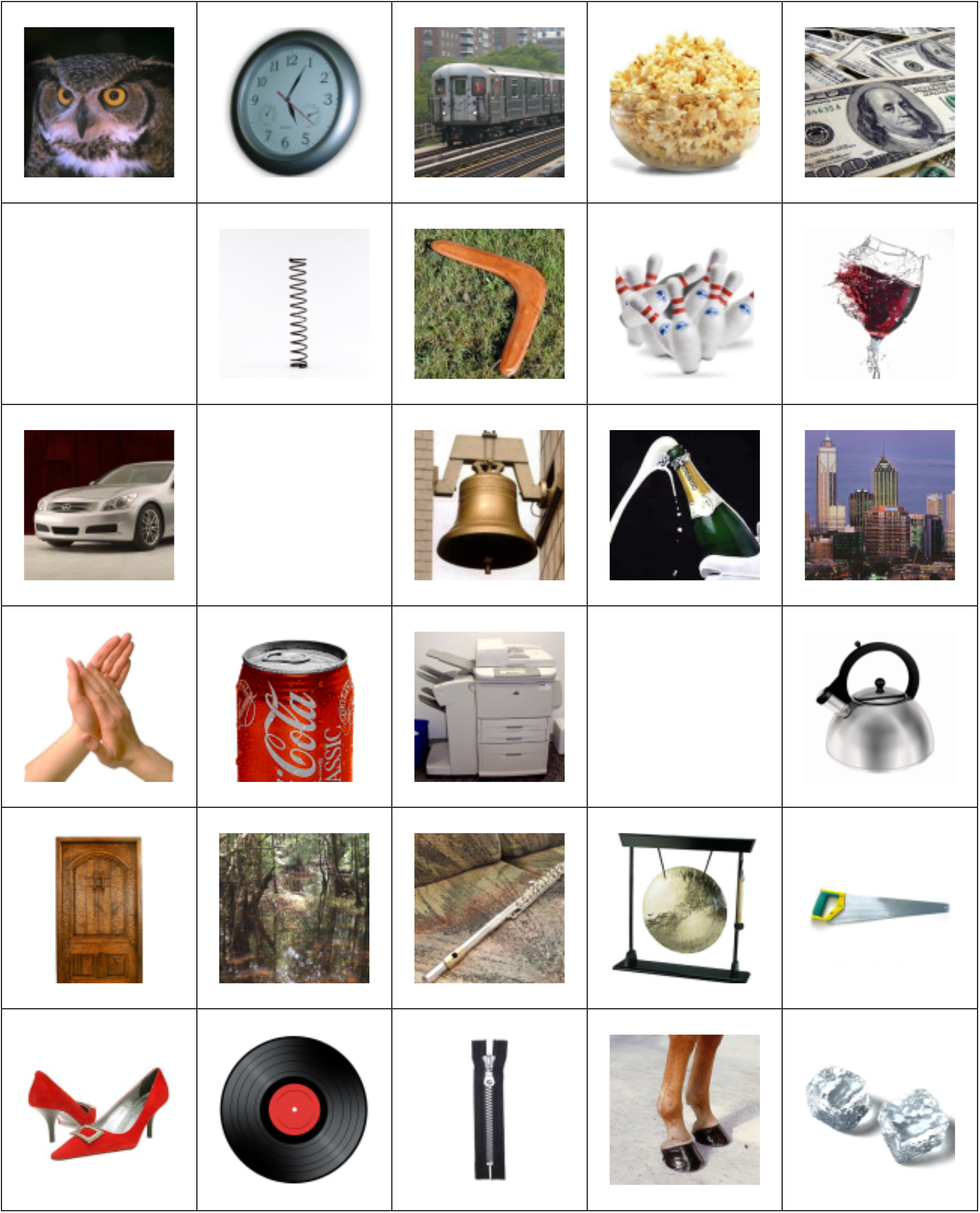
The objects in the memory test. We used these pictures of objects, each with a corresponding sound recording, in a task that required learning object-location associations. For each patient, we adjusted the number of objects in advance by estimating what would tax their abilities to a tolerable degree under the circumstances in the hospital (ranging from 10 to 20, Table 1). Objects were randomly selected from the set of 30 for each patient.

